# Organ-on-a-Chip Fabrication using Dynamic Photomask

**DOI:** 10.1101/2023.11.22.568385

**Authors:** Terry Ching, Shu-Yung Chang, Yi-Chin Toh, Michinao Hashimoto

## Abstract

Organ-on-a-chip (OoC) technology is a powerful tool for creating physiologically relevant microscale models applicable to biomedical studies. Despite the advances in OoC technology, its fabrication method still primarily relies on soft lithography, which (1) lacks the adaptability to accommodate dynamic cell culture (*e.g.*, spheroids and organoid culture) and (2) has a long design-to-prototype cycle that lowers its manufacturability. To overcome these challenges, we developed a system to fabricate OoC (consisting of microchannels and multiple cell types in a well-defined spatial arrangement) dynamically using a digital photomask aligned with a microchamber. Our approach used a pre-defined microfluidic chamber customized by xurography and cell-laden microfluidic channels photopatterned by a digital photomask; the entire design-to-prototype cycle was achieved within two hours. The versatility of our approach offered previously unattainable crucial features in the fabrication of OoC, including a gradual change in the height of the microchannels, and real-time modification of channel designs to trap live tissues (*e.g.*, spheroids). In summary, this work highlights a versatile system to fabricate OoC that can accommodate various design requirements of microenvironments of specific organ tissues. We envision the effectiveness of our system for the rapid fabrication of OoC to contribute to the wide adoption of the technology for therapeutic screening and elucidation of disease mechanisms in both academic and industrial settings.

## 1. Introduction

### Organ-on-a-chip

#### Background

Organ-on-a-chip (OoC), or microphysiological system (MPS), is a miniaturized device designed to recapitulate the structure, microenvironment, and tissue-specific functions of organ tissues. An OoC device achieves physiological relevance to its target tissue *in vivo* in the form of a microfluidic chip that contains cells, extracellular matrix, and perfusion of nutrients and other components essential for the target tissue to function. In the last two decades, research in OoC technology has grown exponentially to model a wide range of organ tissue, including lungs [1–2], heart [3–4], brain [5–6], and liver [7–8]. To capture the interconnectivity of organs *in vivo*, multiple organs were integrated into the chip to emulate the process of therapeutic exposure in humans [9–11]. OoC as an advanced *in vitro* technology has become increasingly important in biomedical research owing to its capability to replicate critical aspects of human physiology, thus providing insights into disease pathophysiology and drug discovery [12–14]. OoC technology has the potential to accelerate clinical research by reducing animal studies and providing more physiologically relevant models. To meet the growing adoption of OoC in the field of biomedical research, it is crucial to improve the process to prototype OoCs, to increase its production efficiency, and to enhance its ability to achieve biomimicry.

#### Fabrication of OoC

Conventional prototyping of OoC devices in research laboratories primarily relies on soft lithography, specifically polydimethylsiloxane (PDMS), which involves multi-step, time-consuming microfabrication [15]. Besides fabrication, the use of PDMS in OoC may result in the adsorption of hydrophobic compounds, leading to complications in therapeutics screening using PDMS-made devices [16]. Additional steps to coat the PDMS surfaces would be required to prevent the adsorption of hydrophobic molecules [16–17]. More importantly, soft lithography does not offer ease of design modifications, as the master mold would have to be fabricated for each device by high-precision photomask printing in a cleanroom [15, 18–20]. This requirement causes a bottleneck in the design-to-prototype iteration (*i.e.,* redesigning the soft lithography mold, fabricating the mold, and fabricating the device) during OoC development. Alternative to PDMS, glass [21] and thermoplastics [22–24] are explored in OoC device fabrication. However, the accessibility to the fabrication tools for the alternative materials (*e.g.*, chemical etching, injection molding, and high-precision laser cutting equipment) has hindered the adoption of these materials in research settings. Recently, additive manufacturing, represented by three-dimensional (3D) printing, of OoC has been explored to enable quick on-site fabrication of OoC. However, the microfluidic devices fabricated by 3D printing remain suboptimal in terms of (1) transparency of the devices, (2) biocompatibility of the available photoresins, and (3) attainable channel dimension and geometry [11].

#### 3D cell culture, spheroids/organoids-on-a-chip

Despite the advances in OoC technologies, challenges remained in reproducing the microenvironment and architecture of the *in vivo* counterparts in these microfluidic systems. The human body is a highly complex entity that comprises organs, each possessing its distinctive microstructure and architecture, complete with its diversity of cellular players [25]. As such, there is a need for a shift from culturing cells as monolayers (*i.e.*, two-dimensional culture) to 3D culture platforms. Besides, the physiological vessels have circular and/or curved cross-sections, which also offer a non-planar arrangement of cells. There have been attempts to incorporate 3D cultures on OoC to recapitulate more complex structures, cell heterogeneity, and functions in OoC. Cell-laden hydrogels have been used to enable the formation of 3D biological tissues within the OoC [12, 26–28]. To further enhance the physiological relevance of OoC, the incorporation of 3D culture in the form of multicellular spheroids (*e.g.*, tumor spheroids and organoids) is necessary. Organoids are a subset of multicellular spheroids with the intrinsic properties to self-organize and develop in a manner resembling organogenesis *in vivo*, making them a powerful tool for the study of disease etiology and therapeutics development. In combination with OoC technology, which allows precise microenvironmental controls, spheroids-on-a-chip (or organoids-on-a-chip) represent a promising *in vitro* technology for therapeutics development and elucidation of disease mechanisms [25, 29–30]. Spheroids-on-a-chip (or organoids-on-a-chip) have led to a need for improved OoC fabrication methods that allow the smooth incorporation of spheroids (or organoids). Current existing systems of spheroids-on-chip often involve the infusion of spheroids suspended in fluids (*e.g.*, culture medium or hydrogel) into the microfluidic channels. The spatial location of the spheroids within the OoC is often uncontrolled [31–33], and this impedes the efficiency of high throughput analysis on microscopy. To immobilize individual spheroids in designated locations, some OoC was designed with micro traps [34–35]. However, the entrapping efficiency of the traps was not very high (*i.e.*, at 75%), and there is a loss of cellular materials that could be scarce, especially in a scenario where cells are patient-derived. One of the strategies to ensure the controlled spatial location of the spheroids without losing them involves the manual transferring of individual spheroids into microwells within the OoC [36–39]. However, this approach is highly user-dependent and time-consuming.

#### Our approach

Overall, despite the progress in the field of OoC, currently available methods to fabricate OoCs are yet to be advanced in terms of (1) time and ease required for design-to-prototype iterations during OoC development, and (2) adaptability to perform post-modification to the fabricated OoCs. To address these limitations, we developed a highly dynamic system to fabricate OoC with a wide variety of designs, while maintaining a design-to-prototype cycle within two hours. This is in contrast with the conventional OoC fabrication with replica molding. In this work, we fabricated OoC with an in-house photocrosslinking instrument consisting of a modified liquid crystal display (LCD) screen that functioned as a digital photomask display. The digital photomask display allowed real-time observation of the OoC fabrication; real-time modification of the photomask to accommodate variability in the shape and sizes of the live tissues (*e.g.*, spheroids and organoids) to be entrapped in the device. With this system, we developed a simple workflow for fabricating OoCs with the following key features: (1) easy customization of microfluidic chip architecture (*i.e.*, inlet and outlets design, height of device, and internal channel architecture) for short design-to-prototype cycles, (2) modification of channel designs during OoC fabrication to accommodate living tissues of varied shapes and sizes. We showed successful endothelialization of tissues on OoC fabricated using our system, where functional endothelial barriers were demonstrated around the cell-laden hydrogels. Our developed system will help to accelerate the innovation of more physiologically relevant OoC devices to meet the growing demand for OoC technologies in both the scientific community and the pharmaceutical industry.

## 2. Materials and Methods

### 2.1 Materials

ibiTreat polymer coverslips were purchased from ibidi (Fitchburg, WI, USA). 3M™ double-coated adhesive tape 9088 was purchased from 3M (3M^TM^, Minnesota, United States). Polyethylene glycol diacrylate (PEGDA) was purchased from Sigma-Aldrich (Missouri, USA). Porcine LunaGel was purchased from Gelomics (Kelvin Grove, QLD, Australia). The BOE 5.5-inch 2160 × 3840 4K monochrome LCD screen (pixel pitch = 31.5 µm) was purchased from Aptus International Co., Limited (Alibaba, Hangzhou, China).

### 2.2 Fabrication of pre-defined microfluidic chambers

The pre-defined microfluidic chamber with the overall structure of the microfluidic chip boundaries was fabricated using two methods. In the first method, it was fabricated by direct ink writing (DIW) of silicone sealant on the coverslip and subsequently sealed with a laser-cut poly(methylmethacrylate) (PMMA) sheet. Details of fabricating the fluidic device are described in our earlier work [40]. The second method was fabricated using adhesive tapes sandwiched between two ibidi polymer coverslips. Briefly, the adhesive tape was cut using a cutting plotter (Silhouette Cameo 2, Silhouette America, Inc., Lindon, UT, USA) to bear the outline of the fluidic chambers. The tape had a thickness of 0.2 mm. Multiple layers of tape were stacked upon each other to increase the thickness of the channel. Holes 1 mm in diameter were created on the polymer coverslips using a biopsy punch to crease inlets and outlets of the chips. The cut adhesive tape was sandwiched between two sheets of polymer coverslips.

### 2.3 Design of photomasks

Photomask was designed in Rhinoceros® (Robert McNeel & Associates, Seattle, USA), and Adobe Illustrator (Adobe, California, USA).

### 2.4 Design of gray-tone photomask

A 3D tubular channels with the desired final dimension was modeled in Rhinoceros® (Robert McNeel & Associates, Seattle, USA). Next, using the Grasshopper® plugin in Rhinoceros®, we mapped out the height of each point on the XY-axis and output a matrix of all the height values where the rows represented the X-axis and the column represented the Y-axis. Using a Python script, we mapped the height values to pixel color values (described in **Fig. 4A**). Lastly, we converted the matrix into a bitmap image file.

### 2.5 Fabrication of channels

To fabricate the channels in the pre-defined microfluidic chamber to form a microfluidic chip, the photomask design for the respective microfluidic chip was displayed on the photocrosslinking instrument to allow photopatterning of the channel features. All photopatterning was performed using ruthenium (RU) and sodium persulfate as photoinitiators at 2 mM and 0.2 mM, respectively. Tartrazine at 0.5 mM was used as a photoabsorber in fabricating tubular channels with a gray-tone mask. Photopatterning of acellular features was performed using 10% (v/v) PEGDA with 6 s exposure to 450 nm light. Photopatterning of cell-laden hydrogels was performed using cells mixed in 1× porcine LunaGel with exposure to 450 nm light for 30 s.

### 2.6 Photopatterning of cell-laden hydrogels for OoC fabrication

Co-culture of RFP-SW480- and GFP-HUVEC-laden hydrogels was performed with 3 million/mL and 8 million/mL of the respective cells in the precursor solution containing LunaGel and photoinitators. Cell-laden hydrogel islands were fabricated by photopatterning 5 million/mL GFP-MCF-7 cells and RFP-MDA-MB-231 cells, respectively, while HUVECs were seeded as monolayers at 5 million/mL after photopatterning of the hydrogel islands.

### 2.7 Immobilization of cancer cell spheroids

To create microwells for cell spheroids formation, molds of the microwell insert for 24-well plates were made using a direct light processing (DLP) 3D printer (Asiga, Alexandria, NSW, Australia). The microwells were made using 14% (v/v) PEGDA with exposure to 450 nm light for 30 s. Subsequently, the microwells were leeched and sterilized by soaking in 70% ethanol overnight to remove uncrosslinked PEGDA monomers. The microwells were then washed and soaked in PBS overnight to remove ethanol residues before the experiment. Each microwell insert contains seven microwells. HepG2 spheroids were pre-formed using 1000 cells per spheroid in the PEGDA microwells. The plate containing the microwells was centrifuged at 1500 rpm for 10 min and placed in the incubator at 37°C and 5% carbon dioxide. The pre-formed spheroids were grown for two days in the microwells before getting harvested for infusion into the pre-defined microfluidic chamber while suspended in LunaGel. For co-culture of HepG2 spheroids with GFP-HUVEC, HepG2 spheroids were pre-formed by seeding 0.3 million cells per well in Aggrewell 800 (Stemcell Technologies, Vancouver, BC, Canada) and centrifuged at 1500 rpm for 10 min before getting transferred into an incubator at 37°C and 5% carbon dioxide. Spheroids were cultured for two days before being suspended in the precursor solution. The spheroids-hydrogel mixture was then infused into the pre-defined microfluidic chamber for photopatterning. A photomask for the photopatterning of GFP-HUVECs was then designed based on the location of the spheroids. Following that, 8 million/mL GFP-HUVEC in precursor solution was then infused into the microfluidic chamber for photopatterning. In the fabrication of hierarchical channels, GFP-HUVEC-laden hydrogels were made with 8 million/mL of cells, while monolayer GFP-HUVEC seeding after the channel formation s was performed with 5 million cells/mL. All cultures were perfused using a 3D-printed peristaltic pump at 0.2 rpm [41]. The spheroids were perfused using a 3D-printed peristaltic pump at 0.2 rpm [41]. Growth of the spheroids was quantified by calculating the area the spheroids occupied when viewed under the microscope (Zeiss, Oberkochen, Germany). Area calculations were performed using FIJI [42].

### 2.8 Evaluation of endothelial barrier

Normal human lung fibroblasts at 3 million/mL and RFP-HUVEC at 8 million/mL in precursor solution were photopatterned to form the endothelial barrier in the microfluidic device generated. The microfluidic chip was perfused with EGM2 medium for 10 days before being used to evaluate barrier integrity using immunofluorescence and 250kDa FITC-dextran perfusion. For FITC-dextran perfusion, microscopy was performed every 15 min, and the relative brightness of the island compared to the channels was quantified on FIJI.

### 2.9 Metastasis on chip

RFP-MDA-MD-231 and GFP-HUVEC at 3 million/mL and 8 million/mL in precursor solution were photopatterned and perfused using the peristaltic pump previously mentioned at 0.2 rpm/mL for five days before tumor necrosis factor-alpha (TNF-α) (R&D Systems, Minneapolis, MN, USA) and Vascular endothelial growth factor (VEGF) (Sigma-Aldrich, St. Louis, MO, USA) were introduced into the channel at 2 µg/mL and 100 µg/mL respectively. Microscopy of the OoC was performed at 0, 3, and 16 h. The number of RFP-MDA-MD-231 cells in the channels was quantified using FIJI.

### 2.10 Immunostaining and confocal microscopy

The microfluidic chips were first washed with 1× PBS twice via perfusion for 5 min each. 4% (wt/v) PFA was then perfused into the channels and incubated for 30 min at room temperature. 1× PBS was then used to rinse the PFA thrice, with 15 min of perfusion per wash. The cells were then permeabilized using 1% (v/v) Triton-X-100 perfusion twice, with 15 min of perfusion per wash at room temperature. Blocking was performed by perfusing blocking buffer, which was composed of 1% (v/v) Triton-X-100, 2% (wt/v) BSA, and 0.2% (wt/v) NaN3 in PBS, 15 min per wash at room temperature thrice. Primary antibodies (anti-VE-Cadherin, 1:200 dilution, catalog # 2500, Cell Signaling Technology, Danvers, MA, USA; anti-ZO-1, 1:100 dilution, catalog # 33-9100, Invitrogen, Waltham, MA, USA) were then perfused into the channels and incubated overnight at 4 °C. The microfluidic chips were then washed using a washing buffer, which is made of 0.2% (v/v) Triton-X-100, 3% (wt/v) NaCl in PBS, thrice at 30 min per wash and a last wash overnight. Secondary antibodies (Alexa Fluor 647 donkey anti-rabbit, catalog # A-31573, Invitrogen; Alexa Fluor 647 donkey anti-mouse, catalog # A-31571, Invitrogen) were then stained at 1:800 dilution together with nuclear stain, SYTOX-green (Invitrogen) at 1:10,000 dilution for 4 – 6 h before being washed in the washing buffer thrice with 30 min per wash and a last overnight wash. Lastly, the channels were washed in PBS thrice before imaging under a ZEISS LSM 700 confocal microscope (ZEISS, Oberkochen, Germany).

### 2.11 Cell culture and maintenance

All cells used in this work were maintained in incubators with 37 °C and 5% carbon dioxide (Thermo Fisher Scientific, Waltham, MA, USA). HUVECs (Lonza, Basel, Switzerland), GFP-HUVECs, and RFP-HUVECs (Angio-Proteomie, Boston, MA, USA) were maintained in EGM-2 endothelial cell growth medium (Lonza, Basel, Switzerland). Cells were passaged at 70 – 80% confluency. Normal human lung fibroblasts (Lonza, Basel, Switzerland) were maintained in FGM-2 Fibroblast growth medium (Lonza, Basel, Switzerland). HepG2, RFP-SW480, GFP-MCF7 (Angio-Proteomie, Boston, MA, USA), and RFP-tagged MDA-MB-231 (GenTarget Inc., San Diego, CA, USA) were maintained in Dulbecco’s Modified Eagle (DMEM) high glucose (Nacalai Tesque, Kyoto, Japan) supplemented with 10% (v/v) fetal bovine serum, 1× Antibiotic-Antimycotic from (Gibco, Waltham, USA) and 0.1% (v/v) MycoZap (Lonza, Basel, Switzerland). Cells were dissociated from the tissue culture flasks (Greiner, Kremsmünster, Austria) using trypsin (Nacalai Tesque, Kyoto, Japan). Cells were incubated with trypsin for 2 – 3 min in a 37 °C incubator. Trypsinized cells were spun down at 1000 rpm for 3 min before getting neutralized using their respective growth media and PBS.

## 3. Results and Discussion

### 3.1 Experimental Design

This research aimed to develop an OoC fabrication system that can be readily available in every laboratory setting and produce OoCs efficiently without the use of PDMS. Conventional OoC fabrication involves the use of soft lithography of PDMS [19] [15, 18, 20] which is resource-intensive and requries long turn-around time. The use of PDMS-based device may not ideal for toxicology and drug screening studies [17]. Our goal is to develop an alternative system— comprising of an instrument and workflow—that facilitate the fabrication of OoCs with flexible design as required by the tissue model of interest.

We investigated to develop a photopatterning system because of (1) short turn-around time (on the order of hours), (2) capability to perform sequential patterning. For the fabrication of the device (containing microchannels and patterned cells), OoCs with a wide variety of biomimetic designs and with resolutions in the micrometer range was successfully fabricated within two hours. The capability for sequntial patterning is particularly crucial to develop OoCs that requires a well-defined spatial arrangement of different cell types. In this work, we demonstrated to pattern *islands* containing different cell types within the same device. The same capability was also demonstrated for post-modification with the microfluidic chamber, which allowed dynamic trapping of spheroids.

We intended to develop a highly adaptable system for OoC fabrication, which was realized using two major components: (1) a customizable microfluidic chamber and (2) a digital photomask display. The microfluidic chambers offered pre-defined areas where the fluidic channels were fabricated. We termed these microfluidic chambers *pre-defined microfluidic chambers*. Stereolithography was used to photopattern hydrogels directly in the chamber for the fabrication of internal microfluidic channel architectures. The customizability of the microfluidic chamber allowed us to control key parameters of the OoC, including the number of inlets and outlets, height, shape, and size of the chamber.

### 3.2 Fabrication of pre-defined microfluidic chambers

In the first step of our fabrication, the pre-defined microfluidic chambers were designed. This microfluidic chamber served as an enclosure with photopatterned microchannels. We fabricated the chamber in two methods: (1) DIW of silicone sealant on the coverslip [40], or (2) xurography [43] (*i.e.*, cutting plot) of 3M double-sided tapes placed on the coverslip (**Fig. 1A (i)**). In either method, the transparency of the coverslip did not compromise the photopatterning of the hydrogels at the subsequent stages. A precursor solution consisting of photocrosslinkable hydrogel was perfused into the pre-defined microfluidic chambers to fill the chamber (**Fig. 1A (ii)**). A digital photomask with the desired pattern was used to project light to crosslink the precursor solution to form the corresponding channel architecture within each chamber (**Fig. 1A (iii)**). The uncured precursor solution was then evacuated by perfusing PBS into the microfluidic chambers (**Fig. 1A (iv)**). Lastly, media reservoirs consisting of PMMA shaped by laser-cutting were attached on top of the chambers to serve as gravity-assisted perfusion. Alternatively, silicone tubes could be attached to the OoC to allow for pump-assisted perfusion (**Fig. 1A (v)**).

**Fig. 1.**
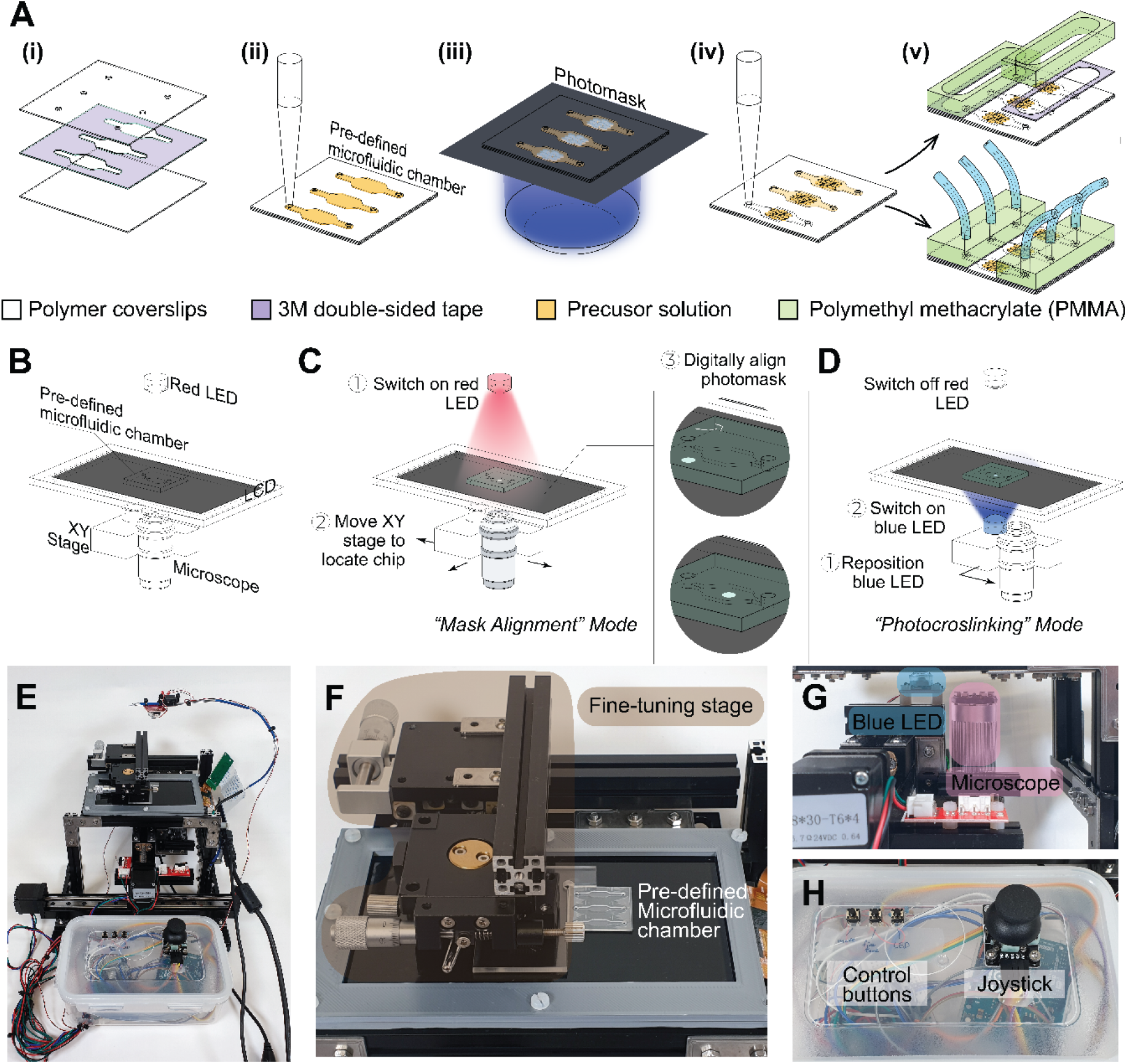
Overview of the system for dynamic OoC fabrication. **(A)** Schematic illustration of the steps (i – v) for the fabrication of OoC using our system. (i) Assembly of polymer coverslips and double-sided tape (purple) after xurography to form the pre-defined microfluidic chamber. (ii) Infusion of precursor solution (orange) consisting of photocrosslinkable hydrogel into the microfluidic chamber. (iii) Photopatterning of microfluidic channels in the chambers using a digital photomask. (iv) Evacuation of uncured precursor from the microfluidic chamber, revealing the microfluidic chip formed. (v) Setting up perfusion systems by attaching media reservoirs made of laser-cut PMMA (green) for gravity-driven perfusion or silicone tubes (blue) for pump-assisted perfusion. **(B)** Schematic illustration showing positions of the red-light emitting diode (LED), pre-defined microfluidic chamber, liquid crystal display (LCD), XY-stage, and microscope on the developed photopatterning instrument. **(C)** Schematic illustration of *Mask Alignment* mode on the photopatterning instrument, in which (1) the red LED was used to illuminate the boundaries of the pre-defined microfluidic chamber for (2) visualization and (3) alignment of the digital photomask with the chamber with the help of the XY-stage. **(D)** Schematic illustration of *Photocrosslinking* mode, in which (1) the blue LED was repositioned to replace the position of the microscope and (2) switched on for photocrosslinking of the precursor solution. **(E)** Photograph showing the overall structure of the developed photopatterning instrument. **(F)** Photograph highlighting the fine-tuning stage (brown) for precise positioning of the microfluidic chambers. **(G)** Photograph highlighting the positions of the blue LED (blue) and the microscope (pink) under the LCD. **(H)** Photograph showing the control box of the photopatterning instrument, where control buttons and joystick were positioned.

### 3.3 Development of photopatterning instrument

We then developed an instrument to perform photopatterning used with the pre-defined microfluidic chamber. This custom-made instrument comprised a monochrome, high-resolution (*i.e.,* pixel size = 32 µm × 32 µm) liquid crystal display (LCD) that functioned as a digital photomask display. The LCD was digitally controlled to switch each pixel on or off selectively. When a pixel was switched on, light was allowed to pass through, and vice versa. The LCD was connected to a computer, which was projecting a live photomask designed by the user. A digital microscope was coupled to a motorized XY-stage below the LCD, which functioned as a living imaging tool to enable the alignment of photomasks with the pre-defined microfluidic chamber (**Fig. 1B**).

To facilitate the operation of the instrument, we designed two modes of operation: *Mask Alignment* and *Photocrosslinking* modes. In *Mask Alignment* mode, the digital microscope was used to align the photomask to the pre-defined microfluidic chamber (**Fig. 1C**). The pre-defined microfluidic chamber was first placed on the LCD screen. Subsequently, a red light-emitting-diode (LED) lamp (wavelength ∼700 nm) directly above the pre-defined microfluidic chamber was illuminated. The red LED light illuminated the boundaries of the pre-defined microfluidic chamber for visualization of the pre-defined microfluidic chamber via the microscope. Red LED light (∼700 nm) was used because it does not interfere with the curing of the photocrosslinkable hydrogels; the photoinitiator used, RU/SPS, had a spectral sensitivity of 400 – 500 nm and was not affected by red light. The motorized XY-stage was then used to maneuver the microscope to locate the pre-defined microfluidic chamber. Then, the mask was digitally controlled to align the photomask with the pre-defined microfluidic chamber. In *Photocrosslinking* mode, the XY-stage automatically positioned a separate 450 nm LED lamp in the position previously occupied by the microscope to photocrosslink the hydrogel in the pre-defined microfluidic chamber (**Fig. 1D**).

The overall structure of the instrument is illustrated (**Fig. 1E**). The digital photomask display (*i.e.*, the LCD) of the instrument was supported by aluminum profiles. To assist in the physical alignment of the pre-defined microfluidic chamber and the photomasks, a fine-tuning stage was incorporated (**Fig. 1F**). The fine-tuning stage allowed fine linear adjustment in the X- and Y-directions and rotation along the Z-axis. Under the LCD screen, the digital microscope and a blue (405 nm) LED lamp were mounted to two electronically controlled linear actuators (**Fig. 1G**). The linear actuator provided movement in the X- and Y-directions. The movements of the linear actuators were controlled by the movement of a joystick on the control box. Three control buttons were used: the first one to toggle between the *mask alignment* and *photocrosslinking* modes, the second one to tune the speed of movement control of the linear actuator, and the last one to switch the blue LED on and off (**Fig. 1H**).

Our instrument enabled two crucial engineering capabilities. First, it allowed rapid customization of the photomask design to perform *in situ* photocrosslinking of hydrogel inside a pre-defined microfluidic chamber. Second, it permitted live imaging of the photomasks coupled with the pre-defined microfluidic chamber; this capability is crucial to properly aligning the pre-defined microfluidic chamber and its features and the photomasks. With the developed instrument, photopatterning of hydrogels could be performed *in situ* within the pre-defined microfluidic chambers to form an OoC that mimics the complex structures of microenvironment (*i.e.,* tissue-level structures, multi-scale organization) found *in vivo*. The live imaging ability of the instrument enabled *in situ* fabrication of channel features, such as the immobilization of dynamic features (such as suspended spheroids) within the patterned microchannel for perfusion culture. These capabilities are discussed in detail in the subsequent sections.

### 3.4 Generating complex 3D biomimetic structures in microfluidic chips

To demonstrate the capability of the instrument, we tested the instrument by photopatterning PEGDA (**Fig. 2A** – **D**). The patterned microchannels suggested that complex networks of interconnecting channels less than 100 µm were achieved (**Fig. 2A, 2B**). The pre-defined microfluidic chambers were readily modified to accommodate the fabrication of dual-channel networks (**Fig. 2C**), and a network mimicking the kidney glomerulus (**Fig. 2D**). Next, to demonstrate the feasibility of our system to create OoC, we fabricated OoC using human cells-laden hydrogels. We encapsulated red fluorescent protein-tagged- (RFP-) SW480 colorectal cancer cells and green fluorescent protein-tagged-(GFP-) HUVEC in gelatin-based hydrogels (**Fig. 2E, 2F**). Crucially, we successfully patterned perfusable channels ∼50 µm in width (**Fig. S1, Supporting Information** and **Fig. 2E**). After five days of continuous perfusion culture, we observed the formation of a sprouts-like structure of the HUVEC (**Fig. 2F**). We observed the sprouting of HUVEC towards the SW480 cells clusters patterned in the core of the hydrogel islands (**Fig. 2F (ii, iii)**). The sprouting of HUVEC towards SW480 cell clusters resembles cancer angiogenesis, where tumor cells secrete angiogenetic factors to induce endothelial sprout formation [39, 44].

**Fig. 2.**
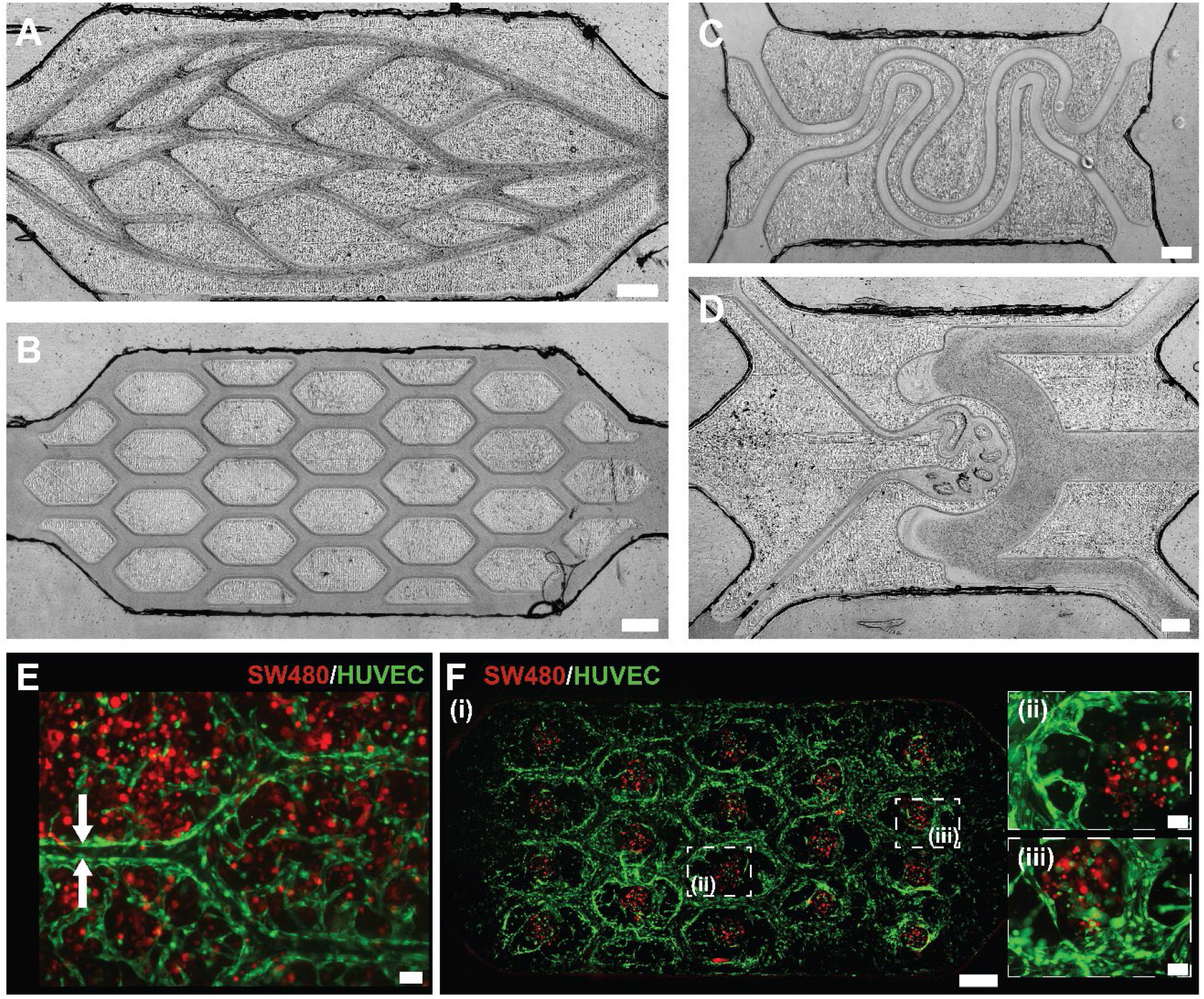
Fabrication of microfluidic chips of varying channel architecture. **(A– D)** Images of microfluidic devices featuring **(A)** interconnecting microchannels mimicking the structure of capillary network complex, **(B)** interconnected microchannels with regular tissue patterns inspired by liver lobules, **(C)** dual inlets/outlets system featuring parallel microchannels mimicking the liver fluid transport system, and **(D)** triple inlet/outlet system featuring channel design inspired by the kidney glomerulus. Scale bars: 500 μm. **(E)** Fluorescence image of RFP-SW480 and GFP-HUVEC in hydrogels perfused by microchannels. White arrows indicate the width of the microchannels fabricated. Scare bar: 100 μm. **(F)** (i) Fluorescence images of RFP-SW480 clustered photopatterned to be surrounded by GFP-HUVEC on a microfluidic chip, and (ii, iii) enlarged views of the dotted sections in (i). Scale bars: (i) 500 μm. (ii, iii) 100 μm.

### 3.5 Fabrication of hierarchical channels

Microchannels fabricated by soft lithography typically offer a uniform height throughout the chip, which poses a limitation in designing a hierarchical channel network with varying widths and heights suitable for some OoCs. This limitation can be readily overcome by the design of the pre-defined chamber in the current work. We devised a pre-defined microfluidic chamber with a gradual change in its cross-sectional height (**Fig. 3A, 3B**). The pre-defined microfluidic chamber was created using adhesive tape of varying heights controlled by xurography. Silicone sealant was utilized to seal the sides of the pre-defined microfluidic chamber between the adhesive tapes of different heights located at the two ends of the chip. This chamber was then used to photocrosslink LunaGel to form a hierarchical branching network with starting channel dimensions at 600 µm (width) by 600 µm (height) that bifurcated twice to form four channels at the end, which had channel dimensions of 150 µm (width) by 100 µm (height). We perfused blue dye to visualize the hierarchical networks and confirmed the formation of the channels (**Movie S1, Supporting Information**).

**Fig. 3.**
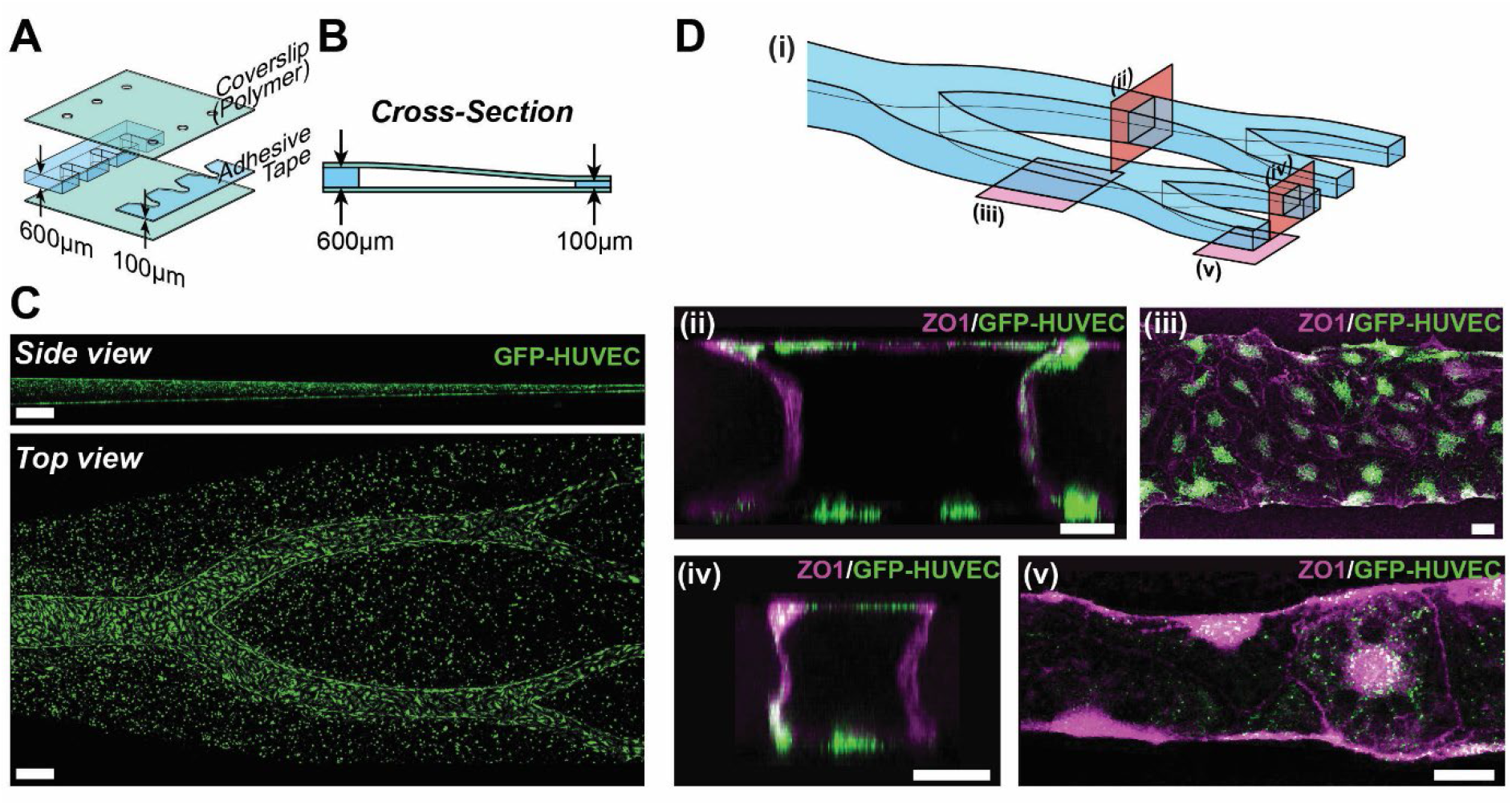
Hierarchical channels with gradual height variation. **(A, B)** Schematic illustration of a pre-defined microfluidic chamber with gradual height change for **(A)** exploded view of the chamber and **(B)** cross-sectional of the chamber. **(C)** Confocal images (side and top views) of hierarchical channels formed by GFP-HUVEC-laden hydrogels featuring the gradual change in both the width and the height of the channels. Scale bars: 500 μm. **(D)** Confocal images of hierarchical channels lined with GFP-HUVEC immunostained with ZO-1 indicating tight junctions. (i) Schematic illustration of the hierarchical channels and the regions of interest. (ii – v) Confocal images of the regions of interest indicated in (i). All scale bars: 50 μm.

**Fig. 4.**
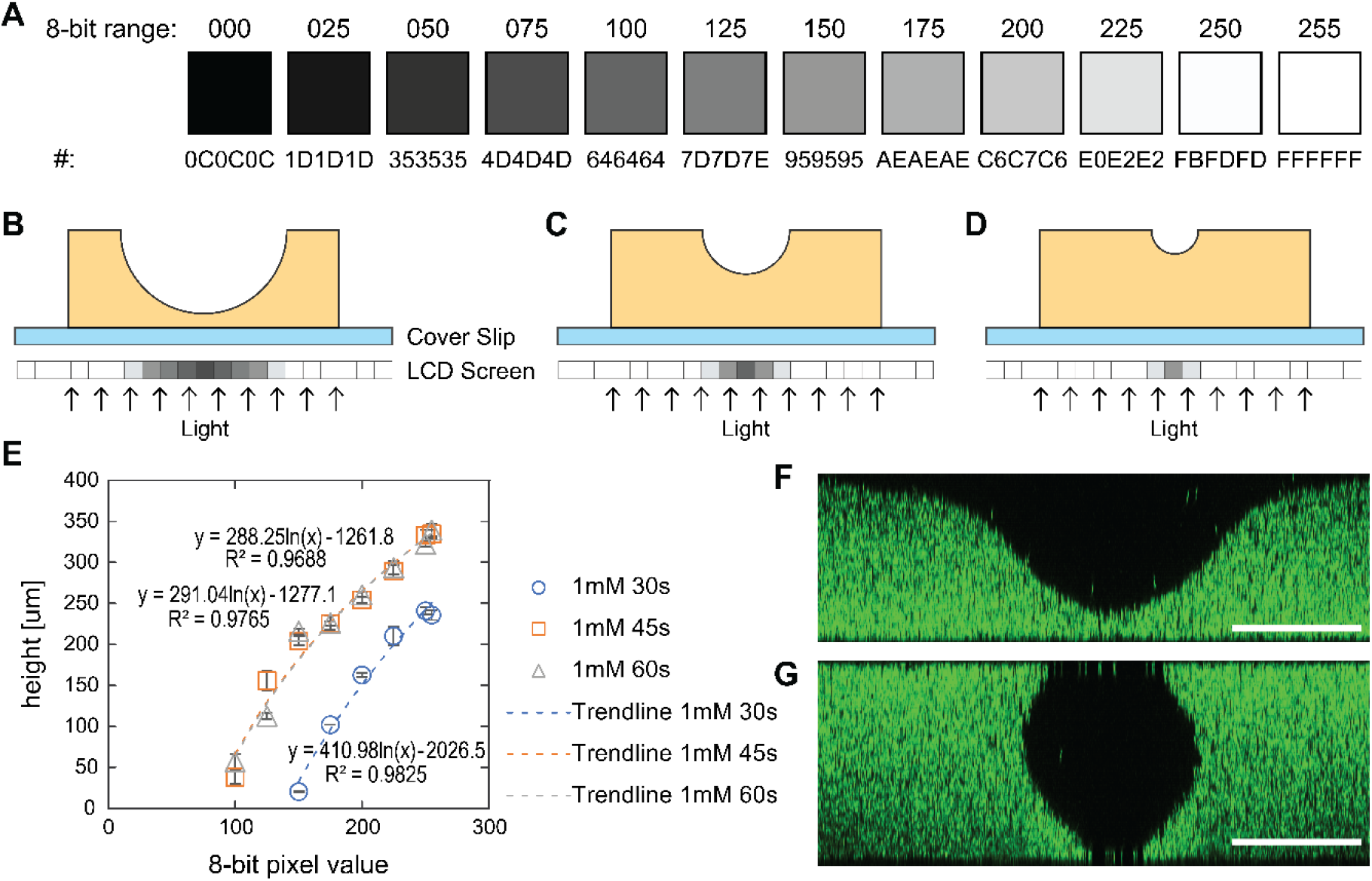
Fabrication of tubular channel. **(A)** Color code corresponding to the 8-bit range (from 000 to 255) employed to control the color of each LCD pixel. **(B – D)** Schematic illustration of the graytone mask for fabricating channels with varying tubular cross-sectional profiles, where (**B**) photomask with more dark pixels creates larger and deeper channel, and decreasing number of dark pixels creates **(C)** smaller and **(D)** shallower channels. **(E)** Graph summarizing the relationship between the height and the 8-bit pixel value (n = 3). **(F, G)** Confocal images of the cross-sectional profile of the hydrogel after **(F)** single exposure on one side and **(G)** exposure on both sides of the chip. Scale bars: 200 μm.

Next, we fabricated the hierarchical network using HUVEC-laden LunaGel. From the maximum projection along the side (*i.e.*, XY-plane) of the hierarchical network fabricated, we observed the gradual decrease in the height of the hierarchical network (**Fig. 3C**). ZO-1 expression showed that the HUVEC formed a confluent layer of cells along the channel walls in the hierarchical network fabricated (**Fig. 3D (ii – v)**). These results demonstrated that our system is capable of fabricating a hierarchical branching network with dimensions not only in the width but also in the height of the channels. The capability to create such channels would contribute to the modeling of various hierarchical fluidic networks in the mesoscopic range (100 – 500 µm) on OoCs. Specifically, modeling the branching of small arteries in the deep cervical arteries, retinal and neocortex regions, and the collecting lymph vessels [45–49].

### 3.6 Graytone Mask

Traditional fabrication of OoC by photolithography does not easily allow for creating channels with circular cross-sections, which is often more physiologically relevant than channels with rectangular cross-sections[50–52]. To this end, we explored the feasibility of using graystone photomasks, enabled by LCD screens, to create tubular channels. An LCD screen as a photomask offers a way to control the transmission of the light through each pixel. The amount of light passing through each pixel could be controlled by the pixel color. This control was readily achieved by changing the 8-bit range from 0 to 255. The corresponding color code for each 8-bit range was summarized (**Fig. 4A**). To create a tubular cross-sectional profile, the pixel color could be varied along the diameter of the desired channel. For the areas where a greater extent of photocrosslinking (in terms of height) is intended, the corresponding pixels on the LCD screen are programmed to transmit an increased amount of light (**Fig. 4B – D**).

We note that the addition of the photoabsorbers to the hydrogels is often required to control and attenuate the light, and the resulting penetration depth of the light. Nordin et al. previously reported the use of photoabsorbers to control the height of photocrosslinked layers [53–54]. In this work, we also used tartrazine as a photoabsorber as it was shown to be biocompatible and non-cytotoxic [55]. This work reported 3D printing of microfluidic devices printed layer-by-layer, where multiple layers (with each layer requiring one-time light exposure) were required to achieve the fabrication of channels with a tubular cross-section. In this work, we achieved the fabrication of the tubular cross-section of channels only with two exposures of light. The measured height of the photocrosslinked hydrogel correlated with the pixel color (*i.e.*, darkness), where an increased transmission of the light resulted in the increased height of the photocrosslinked hydrogel (**Fig. 4E**). We observed a positive correlation between the exposure time and the height of the photocrosslinked hydrogel between 30 s and 45 s. However, increasing the time from 45 s to 60 s did not yield a noticeable difference (**Fig. 4E**). Importantly, the opacity of the pixel and the height of the channel could be correlated to create a graystone mask. Notably, with one single exposure (*e.g.*, 45 s), our method allowed fabricating a semi-circle cross-sectional profile (**Fig. 4F**); the dimension of the channel can be spatially varied using the mask with different opacity. By flipping the microfluidic chip over for the second exposure, channels with a tubular cross-section were achieved (**Fig. 4G**). Overall, this workflow offered a facile way to control channel dimensions and cross-sectional shapes in a microfluidic chip. Existing methods rely mostly on sacrificial molding techniques to create tubular channels in hydrogels [56–57]. Our workflow would offer a rapid and consistent method of fabricating fluidic channels in hydrogels where the alignment of the mask can be digitally controlled.

### 3.7 Spheroids-on-a-chip

Spheoirds-on-a-chip (including organoids-on-a-chip) have been explored extensively to emulate the microarchitecture and functional characteristics of native organs, making them the emerging approach for modeling the development, homeostasis, and disease of various human organs [58]. These spheroids are typically formed (1) in small sizes (< 1 mm diameter), (2) in the form of cellular aggregates suspended in their culture media, and (3) heterogenous in shapes and sizes [59]. This heterogeneity makes it challenging to precisely control the spatial location of the spheroids in microfluidic channels. Without a fixed spatial location allocated for each spheroid, observation, and manipulation of the spheroid experimentally over time is technically difficult. Two-photon polymerization (2PP) has been explored to enable in situ immobilization of tumor spheroids to study cancer cell migration. However, the field of view of the microscope objective necessary for 2PP limits the size of the cages formed and the laser used in this fabrication caused cell damage [60]. This limitation prevents the accommodation of spheroids with increased sizes. Crucially, the commercial instruments of 2PP stereolithography are not readily available due to their cost, and the application of such instruments for exploratory studies in biology is still limited. In contrast, our instrument provided an adaptable way to immobilize in-channel constructs by real-time photopatterning. As our photocrosslinking instrument includes a microscope beneath the digital photomask display, it is possible to observe the microchip in real time. Using the mounted digital microscope, we accurately positioned the digital photomask to confine the spheroids in their respective position and immobilize them for perfusion culture (**Movie S2, Supporting Information**).

To demonstrate the real-time immobilization of spheroids in the microchannel, we first infused the precursor solution containing HepG2 spheroids suspension into our pre-defined microfluidic chamber. Once the spheroids were infused, we aligned the photomask to the exact location of the spheroids. Only the precursor solution exposed to the light (as determined by the photomask) was photocrosslinked to immobilize the spheroids (**Movie S2, Supporting Information**). After immobilizing the spheroids, the fabricated microfluidic chip was used as a perfusion culture of the spheroids at fixed locations (**Fig. 5A**).

**Fig. 5.**
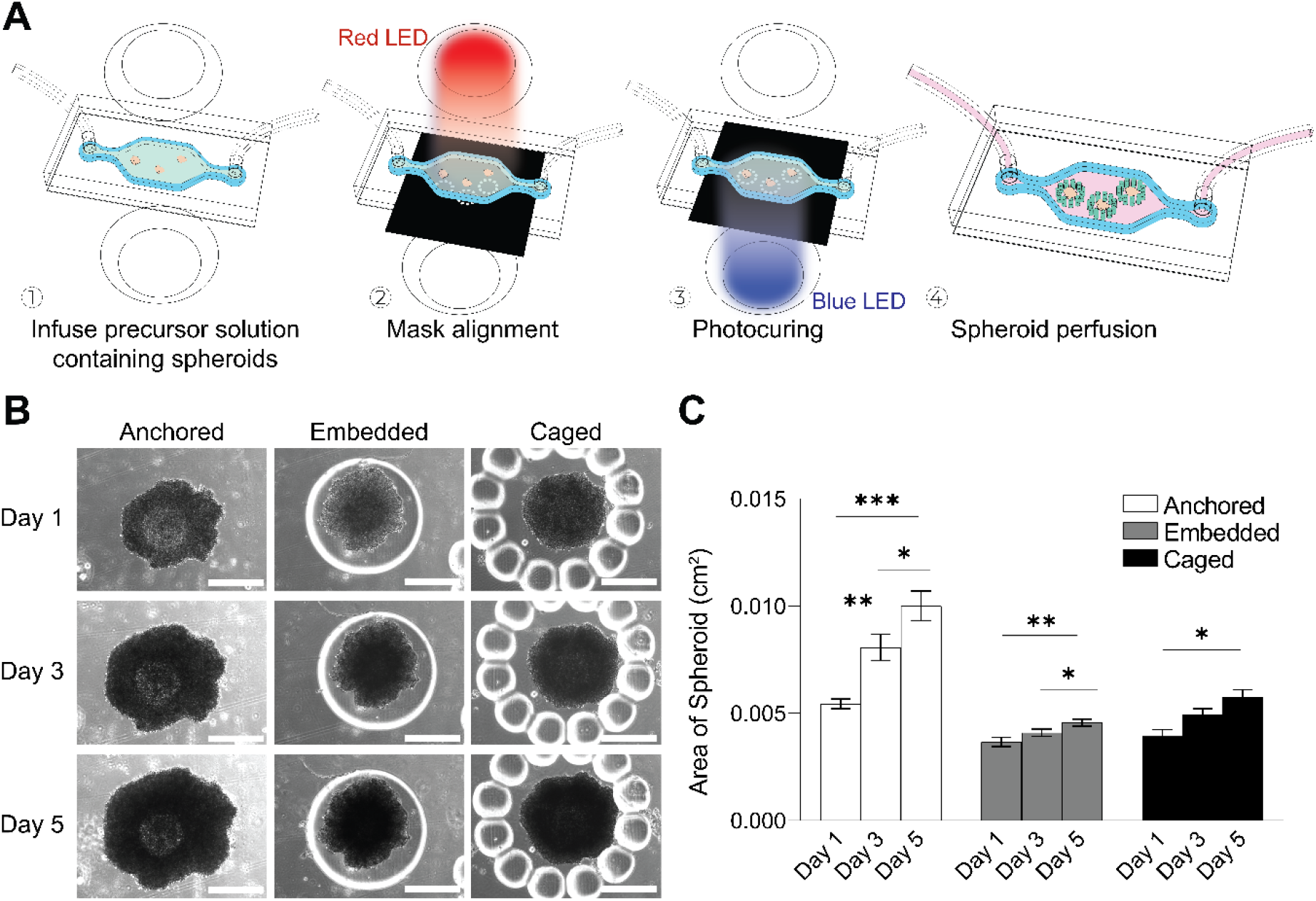
Immobilisation of spheroids-on-chip. **(A)** Schematic illustration of the dynamic workflow to trap HepG2 spheroids in specific locations in the microfluidic chip. **(B)** Microscopic images of the spheroids perfused when anchored, embedded, and caged by micropatterned hydrogels across five days of culture. Scale bars: 500 μm. (C) Quantification of the growth of the spheroids over five days (n = 5). * p < 0.05, ** p < 0.01. *** p < 0.001.

We demonstrated immobilization of the spheroids using three different methods: (1) anchoring the spheroid using a small dot of hydrogel, (2) embedding the entire spheroid in hydrogel, and (3) caging the spheroid in pillars (**Fig. 5B**). This demonstrated the versatility of our system for confining spheroids within a microfluidic chip. We cultured the spheroids for five days under perfusion culture and observed the growth of the spheroids in terms of size across all three methods of immobilization. Among the three methods, anchoring spheroids yielded the highest degree of growth, as seen in the increase in the size of the spheroids observed under bright field microscopy (**Fig. 5C**). This observation could be attributed to the large exposed area the anchored spheroids experienced with the perfusing media in the fluidic chip; in the other two methods, the presence of the hydrogel reduced the flow of the media around the spheroids, resulting in less nutrients transfer. Overall, our system allowed isolating and immobilizing spheroids for perfusion culture for long-term observation under perfusion culture. Crucially, this approach can be readily applied to perform immobilization of organoids for perfusion culture and tissue maturation.

### 3.8 Integrating multiple cell types on a chip

Advantageously, our system allows for the fabrication of OoC with multiple cell types. To this end, islands of hydrogels could be photopatterned successively to create an OoC system with multiple cell types (**Fig. 6A**). The islands within the microfluidic chip could be laden with parenchymal cells, the cells that usually govern the function of organs. Subsequently, the periphery of the islands could be lined with other cell types, such as endothelial cells, to create endothelialized tissues. To demonstrate this OoC system, we fabricated a microfluidic chip with two cancer cell types, MCF-7 human breast cancer (GFP-tagged) and MDA-MB-231 breast cancer cell (RFP-tagged) laden in the hydrogel islands (**Fig. 6B**). The periphery of the islands served as channels, where monolayers of HUVEC were seeded. This successive photopatterning could also be applied to spheroid culture, such that the spheroid in suspension could be immobilized in the first light exposure, followed by photopatterning a second cell type (in this case, GFP-HUVEC) to enable co-culture of cells in two different configurations (*i.e.*, pre-formed 3D spheroid and cells encapsulation in hydrogels) (**Fig. 6C**). This capability to create OoC with multiple cell types and model barrier tissues *in vitro* serves as a powerful tool to predict a multi-organ response to drugs. Alternatively, multiple cancer cell types can be incorporated to gain mechanistic insights and advance our understanding of disease pathophysiology.

**Fig. 6.**
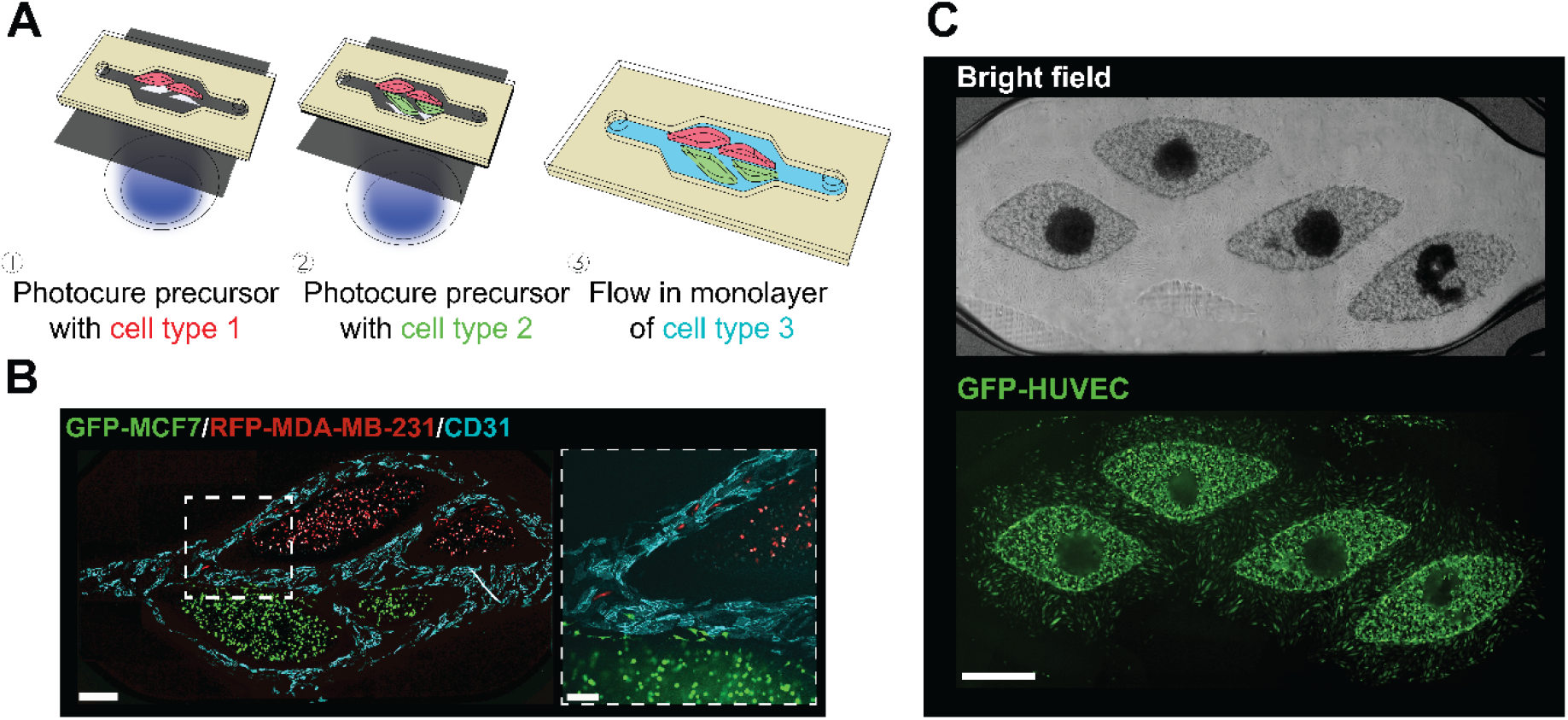
Fabrication of co-culture system using successive exposure. **(A)** Schematic illustration for the steps to fabricate OoC with spatial organization of different cell types using photopatterning. **(B)** Confocal image of spatially organized GFP-MCF7, RFP-MDA-MB-231, and HUVEC immunostained by CD31 on a microfluidic chip. A dotted box showed an enlarged view of a region of interest, highlighting the spatial organization of the cells. Scale bars: 500 μm (right) and 200 μm (left). **(C)** Microscope images of a microfluidic chip with HepG2 spheroids photopatterned in the middle of HUVECs-laden hydrogel. Scale bar: 500 μm.

### 3.9 Characterization of endothelial barriers

Tissues *in vivo* do not interact directly with each other; they interact via vascular flow. The tissues are separated by endothelial barriers, which have selective permeability to molecules and cells so the individual tissues can preserve their specific microenvironments [61]. Ideal multi-organs-on-chip systems offer selectively permeable endothelial barriers between tissues. As such, we characterized the performance of the endothelial barriers fabricated in our method.

To demonstrate the concept of endothelial barriers, we photopatterned precursor solutions containing HUVEC and normal human lung fibroblasts (NHLF) to form islands embedded with the cells. Using immunostaining of VE-Cadherin, we showed that tight junctions were formed between cells in the HUVEC monolayer in our microfluidic chip after eight days of perfusion culture (**Fig. 7A (i)**). From the cross-sectional view of three adjacent hydrogel islands, we confirmed a continuous monolayer of HUVEC lining the walls of the hydrogel islands to form vascularized channels (**Fig. 7A (iii)**). This observation suggested that the tissues grown on OoCs developed using our system could potentially be endothelialized with HUVEC to form endothelial barriers. To further evaluate the barrier function of the HUVEC monolayer grown around the hydrogel islands, we flowed fluorescent dextran (Mw = 250 kDa) into the channels and observed for crossing of the dextran particles into the hydrogel island. Over a period of 60 min, we observed less dextran accumulation in the hydrogel islands surrounded by HUVEC barriers than in the islands without HUVEC barriers (**Fig. 7B – C**). The endothelial barrier formed by the HUVEC monolayer slowed the permeabilization of dextran into the hydrogel island, resulting in reduced dextran accumulation in the hydrogel island with an endothelial barrier. This demonstration suggested the potential to use our approach to fabricate OoC with multiple tissues separated by endothelial barriers through vascular channels and mimic the crosstalk between tissues through vascular systems.

**Fig. 7.**
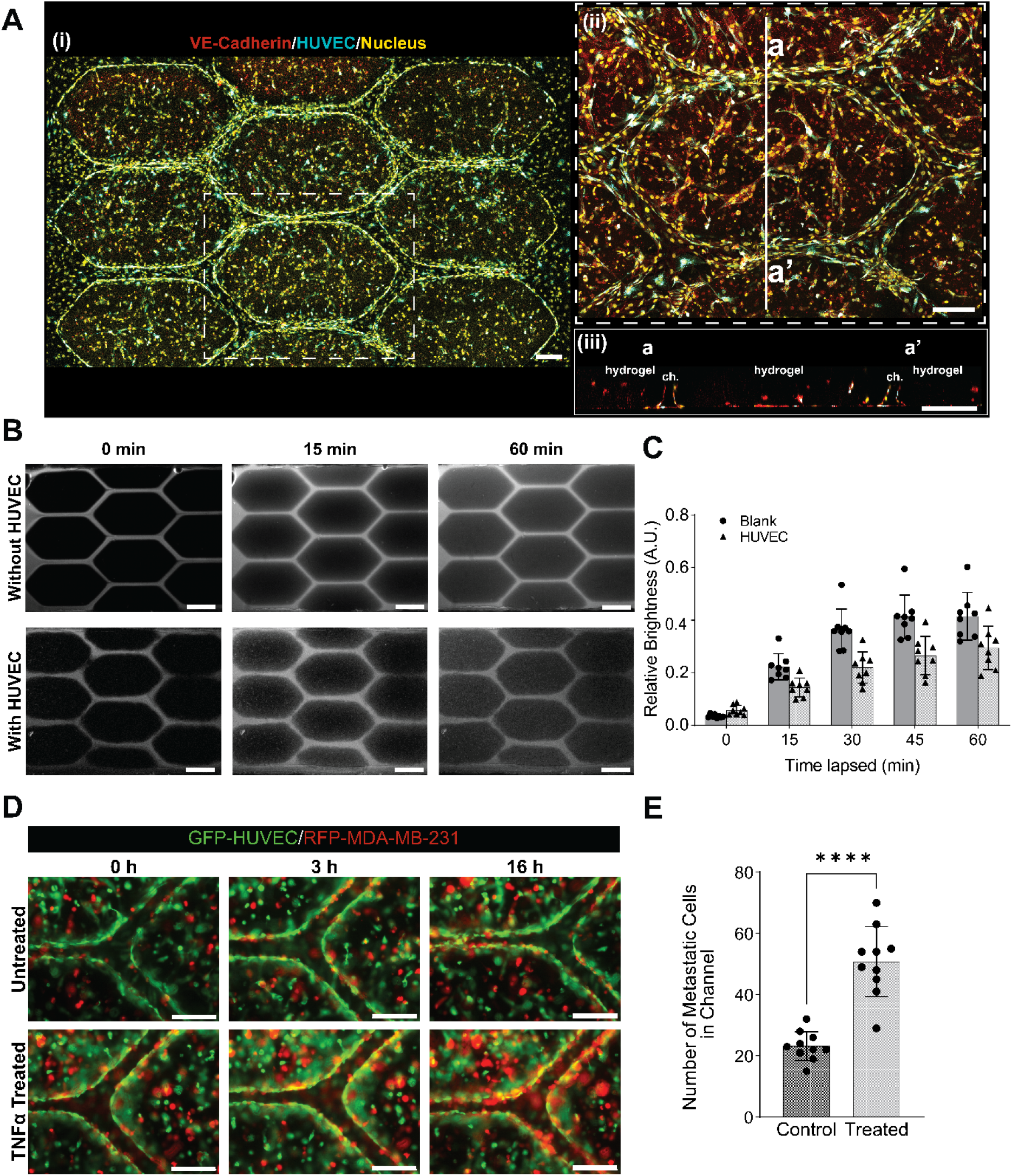
Evaluation of barrier function of the endothelial barrier. **(A)** Confocal images of microfluidic device patterned with hydrogel islands laden by RFP-HUVEC and Normal Human Lung Fibroblasts. VE-Cadherin stains show endothelial barriers formed by RFP-HUVEC around the hydrogel islands: (i) a confocal image, (ii) an enlarged image, and (iii) a cross-sectional image across the line denoted by a – a’. Scale bars: 200 μm. **(B)** Fluorescence images of FITC-dextran (Mw = 250 kDa) in channels at 0, 15, and 60 min after infusion. Scale bars: 500 μm **(C)** Quantification of relative brightness in the hydrogel islands with and without the endothelial barriers formed by HUVECs (n = 8). **(D)** Time-lapse fluorescence images of RFP-MDA-MD-231 and GFP-HUVEC showing migration of metastatic cells through the HUVEC endothelial layer into the channels after treatment with TNF-α. Scale bars: 200 μm. **(E)** Quantification of the number of metastatic cells migrating into the channels 16 h after TNF-α treatment (n = 8), **** p < 0.0001.

To further characterize the functionality of the endothelial barriers formed in our system, we fabricated hydrogel islands using cell-laden hydrogels containing metastatic human breast cancer cells, MDA-MB-231, and HUVEC. To model the migration of metastatic cells through the endothelial barrier, we infused tumor necrosis factor-alpha (TNF-α) through the channels of the microfluidic device. TNF-α has been known to increase endothelial permeability [62]. Over a treatment of 16 hours, we observed MDA-MB-231 cells migrating across the endothelial barrier from the hydrogel islands to populate the endothelial channels (**Fig. 7D**), mimicking the intravasation process, where cancer cells escaped the tumor and invaded the circulatory system [63]. Quantification of the number of MDA-MB-231 cells in the endothelial channels 16 hours after incubation with TNF-α showed that there was a significant amount of metastatic cell migration into the channel (**Fig. 7E**). These demonstrations suggested the potential of our system to create cancer models to study the metastatic processes of the intravasation of cancer cells into the vascular network using vascular-on-a-chip system fabricated using our method.

## 4. Conclusion

In this work, we have developed a versatile system, consisting of a workflow and equipment, to fabricate OoCs with (1) a short design-to-prototype cycle, (2) high adaptability to accommodate the specific design needs of the target organ type, and (3) real-time fabrication of channel structure to interface with dynamic culture types. With highly customizable pre-defined microfluidic chambers made using xurography, our system enabled freedom of design to various parameters of the OoC (*i.e.*, number of inlet and outlets, shape and size of the microfluidic chip, and the height profile of the chip). Our approach can be readily used to fabricate hierarchical channels in both height and width. These features are difficult to fabricate with the conventional fabrication method involving soft lithography. To demonstrate the versatility of channel designs achievable using our system, we employed graytone masks to control the dimension and cross-sectional shape of microchannels in hydrogels, which can be extended to fabricate hierarchical branching networks with circular cross-sectional geometry. Real-time modification of OoC channel designs was demonstrated in the production of spheroids-on-a-chip using our system. The dynamicity of our OoC fabrication allowed real-time immobilization of cancer multicellular spheroids in suspension in different formats for their subsequent perfusion on-chip to encourage their growth. Using cell-laden hydrogels, we demonstrated that our system could be used to fabricate OoCs with 3D spatial organization of multiple cell types and culture formats. Lastly, we showed successful endothelialization of tissues on OoC fabricated using our system, where functional endothelial barriers were formed around the cell-laden hydrogel islands. This enabled us to effectively generate a cancer-on-a-chip to model the intravasation of metastatic cancer cells into the vascular network.

Collectively, we have shown that our system holds great promise to accelerate the progress in OoC technology with the ability to fabricate OoCs with high physiological relevance while reducing the time needed for design-to-prototype iterations to be within two hours. OoC technology has progressed significantly over the last decade with increasing new OoC emerging from academia and the commercialization of OoC devices. However, the adoption of OoC technology in pharmaceutical companies is still in its infancy [64–65]. To accelerate the adoption of OoC technology for routine drug testing and clinical research, there is a need to create a fabrication system that is inexpensive with high usability [12]. We believe that our system contributes to the OoC community as a new tool to rapidly produce a wide range of OoC at a low cost. We believe that the high versatility and usability offered by our system to produce physiologically relevant OoC will lead to an increase in the adoption of OoC in every research setting to enhance the development of new drugs and the understanding of disease etiology.

## Acknowledgements

M.H. acknowledges Agency for Science, Technology and Research (A*STAR) (A*STAR-AMED joint grant, A19B9b0067) and Ministry of Education, Singapore (Academic Research Fund (AcRF) Tier 2, MOE2019-T2-2-192 and MOE-T2EP50122-0025) for the project funding.

## References

[1] D. Huh, B. D. Matthews, A. Mammoto, M. Montoya-Zavala, H. Y. Hsin, D. E. Ingber, Science 2010, 328, 1662.

[2] L. Si, H. Bai, M. Rodas, W. Cao, C. Y. Oh, A. Jiang, R. Moller, D. Hoagland, K. Oishi, S. Horiuchi, S. Uhl, D. Blanco-Melo, R. A. Albrecht, W.-C. Liu, T. Jordan, B. E. Nilsson-Payant, I. Golynker, J. Frere, J. Logue, R. Haupt, M. McGrath, S. Weston, T. Zhang, R. Plebani, M. Soong, A. Nurani, S. M. Kim, D. Y. Zhu, K. H. Benam, G. Goyal, S. E. Gilpin, R. Prantil-Baun, S. P. Gygi, R. K. Powers, K. E. Carlson, M. Frieman, B. R. tenOever, D. E. Ingber, Nat. Biomed. Eng. 2021, DOI: 10.1038/s41551-021-00718-9.

[3] Y. S. Zhang, A. Arneri, S. Bersini, S.-R. Shin, K. Zhu, Z. Goli-Malekabadi, J. Aleman, C. Colosi, F. Busignani, V. Dell’Erba, Biomaterials 2016, 110, 45.

[4] M. Yadid, J. U. Lind, H. A. M. Ardoña, S. P. Sheehy, L. E. Dickinson, F. Eweje, M. M. C. Bastings, B. Pope, B. B. O’Connor, J. R. Straubhaar, B. Budnik, A. G. Kleber, K. K. Parker, Sci. Transl. Med. 2020, 12, eaax8005.

[5] J. Kim, K.-T. Lee, J. S. Lee, J. Shin, B. Cui, K. Yang, Y. S. Choi, N. Choi, S. H. Lee, J.-H. Lee, Y.-S. Bahn, S.-W. Cho, Nat. Biomed. Eng. 2021, 5, 830.

[6] I. Pediaditakis, K. R. Kodella, D. V. Manatakis, C. Y. Le, C. D. Hinojosa, W. Tien-Street, E. S. Manolakos, K. Vekrellis, G. A. Hamilton, L. Ewart, L. L. Rubin, K. Karalis, Nat. Commun. 2021, 12, 5907.

[7] W. L. Chen, C. Edington, E. Suter, J. Yu, J. J. Velazquez, J. G. Velazquez, M. Shockley, E. M. Large, R. Venkataramanan, D. J. Hughes, Biotechnol. Bioeng. 2017, 114, 2648.

[8] N. S. Bhise, V. Manoharan, S. Massa, A. Tamayol, M. Ghaderi, M. Miscuglio, Q. Lang, Y. S. Zhang, S. R. Shin, G. Calzone, Biofabrication 2016, 8, 014101.

[9] I. Maschmeyer, A. K. Lorenz, K. Schimek, T. Hasenberg, A. P. Ramme, J. Hubner, M. Lindner, C. Drewell, S. Bauer, A. Thomas, N. S. Sambo, F. Sonntag, R. Lauster, U. Marx, Lab Chip 2015, 15, 2688.

[10] A. Herland, B. M. Maoz, D. Das, M. R. Somayaji, R. Prantil-Baun, R. Novak, M. Cronce, T. Huffstater, S. S. F. Jeanty, M. Ingram, A. Chalkiadaki, D. Benson Chou, S. Marquez, A. Delahanty, S. Jalili-Firoozinezhad, Y. Milton, A. Sontheimer-Phelps, B. Swenor, O. Levy, K. K. Parker, A. Przekwas, D. E. Ingber, Nat. Biomed. Eng. 2020, 4, 421.

[11] N. Picollet-D’hahan, A. Zuchowska, I. Lemeunier, S. Le Gac, Trends Biotechnol 2021, 39, 788.

[12] T. Ching, Y.-C. Toh, M. Hashimoto, Y. S. Zhang, Trends Pharmacol. Sci. 2021, 42, 715.

[13] S. Cho, S. Lee, S. I. Ahn, Biomed Eng Lett 2023, 13, 97.

[14] D. E. Ingber, Nature Reviews Genetics 2022, 23, 467.

[15] E. Ferrari, F. Nebuloni, M. Rasponi, P. Occhetta, in Organ-on-a-Chip: Methods and Protocols, DOI: 10.1007/978-1-0716-1693-2_1 (Ed: M. Rasponi), Springer US, New York, NY 2022, p. 1.

[16] B. Van Meer, H. De Vries, K. Firth, J. van Weerd, L. Tertoolen, H. Karperien, P. Jonkheijm, C. Denning, A. IJzerman, C. Mummery, Biochem. Biophys. Res. Commun. 2017, 482, 323.

[17] H. Zhang, M. Chiao, J Med Biol Eng 2015, 35, 143.

[18] T. C. Cameron, A. Randhawa, S. M. Grist, T. Bennet, J. Hua, L. G. Alde, T. M. Caffrey, C. L. Wellington, K. C. Cheung, Micromachines (Basel) 2022, 13.

[19] C. M. Leung, P. de Haan, K. Ronaldson-Bouchard, G.-A. Kim, J. Ko, H. S. Rho, Z. Chen, P. Habibovic, N. L. Jeon, S. Takayama, M. L. Shuler, G. Vunjak-Novakovic, O. Frey, E. Verpoorte, Y.-C. Toh, Nature Reviews Methods Primers 2022, 2.

[20] J. Shin, J. Ko, S. Jeong, P. Won, Y. Lee, J. Kim, S. Hong, N. L. Jeon, S. H. Ko, Nat Mater 2021, 20, 100.

[21] H. Hirama, T. Satoh, S. Sugiura, K. Shin, R. Onuki-Nagasaki, T. Kanamori, T. Inoue, Journal of Bioscience and Bioengineering 2019, 127, 641.

[22] S. Kim, J. Ko, S. R. Lee, D. Park, S. Park, N. L. Jeon, Biotechnol Bioeng 2021, 118, 2524.

[23] M. Gröger, J. Dinger, M. Kiehntopf, F. T. Peters, U. Rauen, A. S. Mosig, Adv. Healthc. Mater. 2018, 7, 1700616.

[24] S. A. M. Shaegh, A. Pourmand, M. Nabavinia, H. Avci, A. Tamayol, P. Mostafalu, H. B. Ghavifekr, E. N. Aghdam, M. R. Dokmeci, A. Khademhosseini, Y. S. Zhang, Sens. Actuators B Chem. 2018, 255, 100.

[25] D. R. Mertz, T. Ahmed, S. Takayama, Lab Chip 2018, 18, 2378.

[26] J. E. Ortiz-Cárdenas, J. M. Zatorski, A. Arneja, A. N. Montalbine, J. M. Munson, C. J. Luckey, R. R. Pompano, Organs-on-a-Chip 2022, 4, 100018.

[27] J. Paek, S. E. Park, Q. Lu, K.-T. Park, M. Cho, J. M. Oh, K. W. Kwon, Y.-s. Yi, J. W. Song, H. I. Edelstein, J. Ishibashi, W. Yang, J. W. Myerson, R. Y. Kiseleva, P. Aprelev, E. D. Hood, D. Stambolian, P. Seale, V. R. Muzykantov, D. Huh, ACS Nano 2019, 13, 7627.

[28] V. S. Shirure, Y. Bi, M. B. Curtis, A. Lezia, M. M. Goedegebuure, S. P. Goedegebuure, R. Aft, R. C. Fields, S. C. George, Lab Chip 2018, 18, 3687.

[29] G. Fang, Y. C. Chen, H. Lu, D. Jin, Adv. Funct. Mater. 2023, 33.

[30] T. Takebe, B. Zhang, M. Radisic, Cell Stem Cell 2017, 21, 297.

[31] Y. Wang, L. Wang, Y. Zhu, J. Qin, Lab Chip 2018, 18, 851.

[32] K. A. Homan, N. Gupta, K. T. Kroll, D. B. Kolesky, M. Skylar-Scott, T. Miyoshi, D. Mau, M. T. Valerius, T. Ferrante, J. V. Bonventre, J. A. Lewis, R. Morizane, Nat Methods 2019, 16, 255.

[33] K. Chen, M. Wu, F. Guo, P. Li, C. Y. Chan, Z. Mao, S. Li, L. Ren, R. Zhang, T. J. Huang, Lab Chip 2016, 16, 2636.

[34] B.-J. Jin, S. Battula, N. Zachos, O. Kovbasnjuk, J. Fawlke-Abel, J. In, M. Donowitz, A. S. Verkman, Biomicrofluidics 2014, 8.

[35] A. Schepers, C. Li, A. Chhabra, B. T. Seney, S. Bhatia, Lab Chip 2016, 16, 2644.

[36] K. K. Lee, H. A. McCauley, T. R. Broda, M. J. Kofron, J. M. Wells, C. I. Hong, Lab Chip 2018, 18, 3079.

[37] S. Rajasekar, D. S. Y. Lin, L. Abdul, A. Liu, A. Sotra, F. Zhang, B. Zhang, Adv Mater 2020, 32, e2002974.

[38] Z. Hu, Y. Cao, E. A. Galan, L. Hao, H. Zhao, J. Tang, G. Sang, H. Wang, B. Xu, S. Ma, ACS Biomater Sci Eng 2022, 8, 1215.

[39] J. Ko, J. Ahn, S. Kim, Y. Lee, J. Lee, D. Park, N. L. Jeon, Lab Chip 2019, 19, 2822.

[40] T. Ching, Y. Li, R. Karyappa, A. Ohno, Y.-C. Toh, M. Hashimoto, Sens. Actuators B Chem. 2019, 297, 126609.

[41] T. Ching, J. Vasudevan, H. Y. Tan, C. T. Lim, J. Fernandez, Y. C. Toh, M. Hashimoto, HardwareX 2021, 10, e00202.

[42] J. Schindelin, I. Arganda-Carreras, E. Frise, V. Kaynig, M. Longair, T. Pietzsch, S. Preibisch, C. Rueden, S. Saalfeld, B. Schmid, J. Y. Tinevez, D. J. White, V. Hartenstein, K. Eliceiri, P. Tomancak, A. Cardona, Nat Methods 2012, 9, 676.

[43] T. Hizawa, A. Takano, P. Parthiban, P. S. Doyle, E. Iwase, M. Hashimoto, Biomicrofluidics 2018, 12.

[44] S. Lee, S. Kim, D. J. Koo, J. Yu, H. Cho, H. Lee, J. M. Song, S. Y. Kim, D. H. Min, N. L. Jeon, ACS Nano 2021, 15, 338.

[45] M. Arslan, H. I. Acar, A. Comert, R. S. Tubbs, Turk Neurosurg 2018, 28, 234.

[46] T. H. Rim, Y. S. Choi, S. S. Kim, M. J. Kang, J. Oh, S. Park, S. H. Byeon, Eye (Lond) 2016, 30, 111.

[47] S. Bollmann, H. Mattern, M. Bernier, S. D. Robinson, D. Park, O. Speck, J. R. Polimeni, Elife 2022, 11.

[48] H. Hara, M. Mihara, J Vasc Surg Venous Lymphat Disord 2022, 10, 758.

[49] W. R. Pan, C. M. le Roux, S. M. Levy, C. A. Briggs, Clin Anat 2010, 23, 654.

[50] J. P. Camp, T. Stokol, M. L. Shuler, Biomed Microdevices 2008, 10, 179.

[51] A. Pollet, E. Homburg, R. Cardinaels, J. M. J. den Toonder, Micromachines (Basel) 2019, 11.

[52] S.-H. Song, C.-K. Lee, T.-J. Kim, I.-c. Shin, S.-C. Jun, H.-I. Jung, Microfluid. Nanofluidics 2010, 9, 533.

[53] H. Gong, B. P. Bickham, A. T. Woolley, G. P. Nordin, Lab Chip 2017, 17, 2899.

[54] H. Gong, M. Beauchamp, S. Perry, A. T. Woolley, G. P. Nordin, RSC Adv. 2015, 5, 106621.

[55] B. Grigoryan, S. J. Paulsen, D. C. Corbett, D. W. Sazer, C. L. Fortin, A. J. Zaita, P. T. Greenfield, N. J. Calafat, J. P. Gounley, A. H. Ta, Science 2019, 364, 458.

[56] H. H. G. Song, A. Lammers, S. Sundaram, L. Rubio, A. X. Chen, L. Li, J. Eyckmans, S. N. Bhatia, C. S. Chen, Adv. Funct. Mater. 2020, 30, 2003777.

[57] I. S. Kinstlinger, S. H. Saxton, G. A. Calderon, K. V. Ruiz, D. R. Yalacki, P. R. Deme, J. E. Rosenkrantz, J. D. Louis-Rosenberg, F. Johansson, K. D. Janson, Nat. Biomed. Eng. 2020, 4, 916.

[58] S. E. Park, A. Georgescu, D. Huh, Science 2019, 364, 960.

[59] M. A. Lancaster, N. S. Corsini, S. Wolfinger, E. H. Gustafson, A. W. Phillips, T. R. Burkard, T. Otani, F. J. Livesey, J. A. Knoblich, Nat. Biotechnol. 2017, 35, 659.

[60] M. Taale, B. Schamberger, F. Taheri, Y. Antonelli, A. Leal-Egaña, C. Selhuber-Unkel, Adv. Funct. Mater. 2023, DOI: 10.1002/adfm.202302356.

[61] K. Ronaldson-Bouchard, D. Teles, K. Yeager, D. N. Tavakol, Y. Zhao, A. Chramiec, S. Tagore, M. Summers, S. Stylianos, M. Tamargo, B. M. Lee, S. P. Halligan, E. H. Abaci, Z. Guo, J. Jackow, A. Pappalardo, J. Shih, R. K. Soni, S. Sonar, C. German, A. M. Christiano, A. Califano, K. K. Hirschi, C. S. Chen, A. Przekwas, G. Vunjak-Novakovic, Nat Biomed Eng 2022, 6, 351.

[62] J. Friedl, M. Puhlmann, D. L. Bartlett, S. K. Libutti, E. N. Turner, M. F. X. Gnant, H. R. Alexander, Blood 2002, 100, 1334.

[63] A. F. Chambers, A. C. Groom, I. C. MacDonald, Nat Rev Cancer 2002, 2, 563.

[64] K. A. Homan, Adv Biol (Weinh) 2023, 7, e2200334.

[65] P. Vulto, J. Joore, Nat Rev Drug Discov 2021, 20, 961.

