## Supplementary Information for "Organ-on-a-Chip Fabrication using Dynamic Photomask"

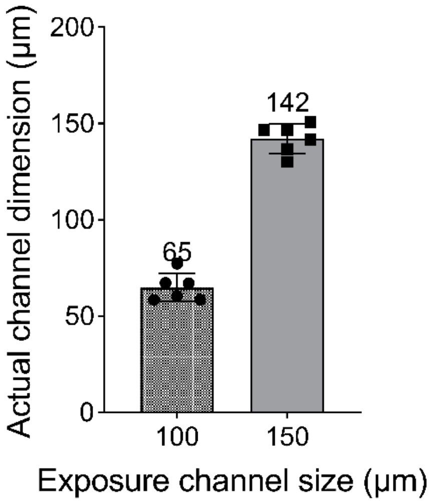


**Fig. S1. Photopatterned channel dimensions.** Quantification of the channel width against the designed channel width (n = 6).
